# Enhanced sensitivity of TAPscan v4 enables comprehensive analysis of streptophyte transcription factor evolution

**DOI:** 10.1101/2024.07.13.602682

**Authors:** Romy Petroll, Deepti Varshney, Saskia Hiltemann, Hermann Finke, Mona Schreiber, Jan de Vries, Stefan A. Rensing

## Abstract

Transcription associated proteins (TAPs) fulfill multiple functions in regulatory and developmental processes and display lineage-specific evolution. TAPscan is a comprehensive and highly reliable tool for genome-wide TAP annotation via domain profiles. Here, we present TAPscan v4, including an updated web interface (https://tapscan.plantcode.cup.uni-freiburg.de/), which enables an in-depth representation of the distribution of 138 TAP families across 678 species from diverse groups of organisms, with a focus on Archaeplastida (plants in the wide sense). With this release, we also make the underlying “Genome Zoo” available, a curated protein data set with scripts and metadata. 18 new TAP (sub)families were added as part of the update. Nine of those were gained in the most recent common ancestor of the Streptophyta (comprising streptophyte algae and land plants), or within the streptophyte algae. More than one-third of all detected TAP family gains were identified during the evolution of streptophyte algae, before the emergence of land plants, and are thus likely to have been significant for plant terrestrialization. The TAP complement of the Zygnematophyceae was identified to be the most similar to that of land plants, consistent with the finding that this lineage is sister to land plants. Overall, our data retrace the evolution of streptophyte TAPs, allowing us to pinpoint the regulatory repertoire of the earliest land plants.

## 1. Introduction

Transcription associated proteins (TAPs) are involved in transcriptional regulation and thus essential players in gene regulatory networks (GRNs). TAPs can be classified into transcription factors (TFs) and transcriptional regulators (TRs). TFs bind sequence-specifically to regulatory elements, resulting in enhancing or repressing of transcription (Richardt et al., 2007; Wilhelmsson et al., 2017). In contrast, TRs are involved in protein-protein interactions, may serve as regulators at the transcriptional core complex, as co-activators and co-repressors, or can be involved in chromatin modification or methylation (Richardt et al., 2007; Wilhelmsson et al., 2017). Furthermore, there are proteins referred to as putative TAPs (PTs) that are thought to be involved in the regulation of transcription, but their exact function is undefined (Richardt et al., 2007). The TAPscan tool has been established for comprehensive genome-wide annotation of TAP families using HMM profiles (Richardt et al., 2007; Wilhelmsson et al., 2017). While the main focus originally was plants, it has recently been improved for detection in several lineages of algae (Petroll et al., 2021).

A possible definition of the morphological complexity of an organism is the complexity of GRNs and their underlying TAP complement (Lang & Rensing, 2015). Within the Chloroplastida, the green lineage (Adl et al., 2012), a correlation between an increase in the complexity of the TAP complement of an organism and its morphological complexity — defined by the number of different cell types — was suggested (Lang et al., 2010; Lang & Rensing, 2015). The Chloroplastida are part of the Archaeplastida, which also comprise the Rhodophyta (red algae) and Glaucophyta (Leebens-Mack et al., 2019). The Chloroplastida include the Chlorophyta (green algae), the Prasinodermophyta, and the Streptophyta (Becker & Marin, 2009; L. Li et al., 2020). The Streptophyta are a monophyletic clade that includes the streptophyte algae and the Embryophyta, the land plants (Leebens-Mack et al., 2019). The paraphyletic grade of streptophyte algae comprises six distinct major lineages, namely Klebsormidiophyceae, Chlorokybophyceae, and Mesostigmatophyceae (summarized as KCM grade), and Zygnematophyceae, Charophyceae, and Coleochaetophyceae (summarized as ZCC grade) (J. de Vries et al., 2016). The ZCC grade together with the Embryophyta forms the monophyletic Phragmoplastophyta, united by the presence of traits such as polyplastidy and the phragmoplast (Buschmann & Zachgo, 2016; Nishiyama et al., 2018).

The archaeplastidal ancestor and its successors dwelled in aquatic environments (J. de Vries et al., 2016) — likely in freshwater (Sánchez-Baracaldo et al., 2017). While several lineages of algae have made a water-to-land transition, the event about 500 Ma ago was unique (Fürst-Jansen et al., 2020): the water-to-land transition that gave rise to embryophytes (land plants), also referred to as plant terrestrialization (Rensing, 2018). Through phylogenetic analyses it was demonstrated that land plants and Zygnematophyceae are sister lineages (Cheng et al., 2019; Leebens-Mack et al., 2019). Hence, the conquest of land was most likely accomplished by the common ancestor of these freshwater streptophyte algae and the Embryophyta (J. de Vries et al., 2016; J. de Vries & Archibald, 2018). Until recently, Zygnematophyceae were not considered the closest relatives to land plants, since their extant species display the lowest morphological complexity within the ZCC grade and are therefore morphologically least similar to their sister group, the embryophytes (J. de Vries & Archibald, 2018). Zygnematophyceae are the richest in species numbers of all streptophyte algae and comprise unicellular and simple filamentous species spread out over five orders (Hess et al., 2022) but lack morphological adaptations like apical growth or the occurrence of plasmodesmata (Cheng et al., 2019; Moody, 2020). In contrast, Charophyceae, earlier thought to be the closest land plant relatives, have multicellular land plant-like body plans with a shoot-like axis, branched nodes, apical meristem, and rhizoids (Nishiyama et al., 2018). Therefore, the simple body plans of extant Zygnematophyceae suggests a reductive evolution of morphological complexity within this lineage (J. de Vries & Archibald, 2018; Hess et al., 2022; Wickett et al., 2014; Wodniok et al., 2011). In general, it needs to be considered that it remains unknown how exactly the ancestor looked like, and which features have emerged or been lost during evolution (J. de Vries & Archibald, 2018). For this reason, it is essential to study extant freshwater relatives of land plants together with the bryophytes, sister clade to all other land plants, in order to infer about the process of terrestrialization (Rensing, 2018).

Terrestrial life faces many challenges, especially due to abiotic stresses like drought, temperature changes, and high-intensity light radiation. Therefore, specific adaptations were required for life on land. Aquatic streptophyte algae are freshwater algae and thus potentially better adapted to conditions on land than marine algae, since the freshwater environment is entangled with the environment on land (Delwiche & Cooper, 2015). Of note, besides the aquatic species within the streptophyte algae, there are also those that live in subaerial/terrestrial environments, which dwell on wet soil or on rock surfaces (Bierenbroodspot et al., 2024; J. de Vries & Archibald, 2018; Fürst-Jansen et al., 2020; Karsten & Holzinger, 2014; Lewis & McCourt, 2004; McCourt et al., 2023). Also, streptophyte algae possess multiple beneficial features for life on land, such as the ability to resist desiccation, flexible cell walls, an expanded repertoire of phenolics to protect against UV damage— possibly from the phenylpropanoid pathway, among others (J. de Vries & Archibald, 2018; S. de Vries et al., 2021; Herburger & Holzinger, 2015; Karsten & Holzinger, 2014; Remias et al., 2012; Rieseberg et al., 2023). Based on their ability to deal with drought, species belonging to the Zygnematophyceae are summarized together with the land plants as Anydrophyta, plants that can resist drought (Rensing, 2020). In conclusion, the ancestor of the Phragmoplastophyta probably already possessed features that enabled it to adapt to the terrestrial environment, and further adaptations allowed a broad diversity of land plants to arise.

Here, we performed an analysis of the TAP complement of Archaeplastida, making use of several genomes of recently sequenced streptophyte algae. We introduce a comprehensive update of the tool TAPscan to v4, enabling to detect 18 additional TAP (sub)families. We also included organismal groups outside the Archaeplastida for comparative purposes.

## 2. Materials and Methods

### 2.1 Data sets

Three data sets (Table S1) were used for different steps of analyses: data set 1 (DS1) is a species-diverse dataset used for determining HMM cutoffs for newly added (sub)families, DS2 is a dataset that focuses on streptophyte algae, for the purpose of gaining new insights into early TAP evolution in Streptophyta, while DS3 is a broad species representation used for the v4 web interface. An overview of the data sets can be found in Table S1.

The data sets DS1 and DS2 include published genome data of species belonging to the Archaeplastida as well as additional species belonging to SAR-group (Stramenopiles, Alveolates, and Rhizaria) members (since they comprise photosynthetic eukaryotes such as brown algae). DS1 was used to add new TAP families and to determine thresholds. It consists of a diverse group of 42 species including the genomes of two Cryptophyta (used as outgroup), four Rhodophyta, one Glaucophyta, five Chlorophyta, two streptophyte algae belonging to the KCM grade, four streptophyte algae of the ZCC grade, one hornwort, one liverwort, three mosses, one lycophyte, one fern, two Acrogymnospermae, one species belonging to the ANA grade, five eudicots, three monocots (Liliopsida), and six SAR-group members. This selection formed the sequence basis for adding new TAP families, to subsequently allow identification of these TAP families in a broad range of species with a focus on the Archaeplastida.

DS2 was used for the analysis of the TAP complements in streptophyte algae and differed from DS1 in the sense that no SAR group members are included and a reduced number of species belonging to the embryophytes were chosen (in total 16) to adjust the proportion of the number of embryophytes as compared to streptophyte algae (Table S1). Also, two further recently sequenced species belonging to the streptophyte algae and one species belonging to the recently identified Prasinodermophyta were added. Thus, the data set consisted in total of 37 species, with 30 species belonging to the Chloroplastida (green lineage). The complete annotated TAPs of these species using TAPscan v4 can be found in Table S2. In addition, for the principal component analysis (PCA), only 23 of the 30 species belonging to the Chloroplastida were used, with fewer embryophytes utilized for brevity.

The analysis conducted for the new TAPscan v4 webpage utilized a broad set of species from MAdLandDB/Genome Zoo (DS3). The Genome Zoo has been expanded in the framework of the priority program MAdLand (https://madland.science/) and served as the basis for previous TAPscan versions. It contains over 21 million representative protein isoforms from a diverse group of nearly 700 fully sequenced species including plants, algae, fungi, mammals, SAR group members, bacteria, and archaea (Table S1). The data are primarily from genome projects and make use of isoform information if present, to represent each locus by its representative isoform only, reducing redundancy. Python scripts are used to refine and extract the representative protein sequences for each species. After downloading the protein data set from the source repository, we use a standardized species abbreviation method to incorporate a leading five-letter code derived from the first three letters of the genus and the first two letters of the species name (e.g., ORYSA = ORYza SAtiva) in the sequence header. This code allows straightforward identification of species and at the same time keeps the original (tailing) identifier. We also include information about the gene’s encoding source via a two character suffix following the five-letter code if the source is not the nuclear genome (pt=plastid, mt=mitochondrion or pl=plasmid). Further suffix information is encoded by specific abbreviations, namely tr for transcriptome, hc for high-confidence proteins, lc for low-confidence proteins, and org for organellar proteins (applied when the origin of the organellar proteins is ambiguous). In versions of the data set that include isoform information, iso is used to denote non-major isoform proteins. The MAdLandDB/Genome Zoo dataset is available via the Galaxy platform (Afgan et al., 2022), allowing for NCBI BLAST and Diamond searches for comparative analyses. All the protein fasta files, source and metadata as well as scripts are available on https://github.com/Rensing-Lab/Genome-Zoo.

### 2.2 Procedure to add new TAP (sub)families

To decide which TAP families to add or sub-divide to improve classification, we used recent publications as gold standards for each revised TAP family. These publications included investigations of the plant model organism *Arabidopsis thaliana* in terms of domain analyses and/or phylogenetic analyses. An exception is the subfamily HD_TALE, for which the brown alga *Ectocarpus siliculosus* was used as gold standard, to ensure detection beyond Archaeplastida. The gold standards have been collected by performing PubMed query searches and manual literature searches (Sayers et al., 2021). As the first step of adding new TAP families to TAPscan and since the annotation of TAPs is based on the classification of protein domains, based on the selected gold standard publications a profile HMM for each protein domain was retrieved. To ensure high sensitivity and specificity of these domains, the second step was the calculation of specific cutoffs, thresholds and classification rules as previously introduced (Wilhelmsson et al., 2017). A summary of which gold standards were used for each new TAP (sub)family can be found in Table S3. The procedure to add new TAP (sub)families is shown in Figure 1.

**Figure 1.**
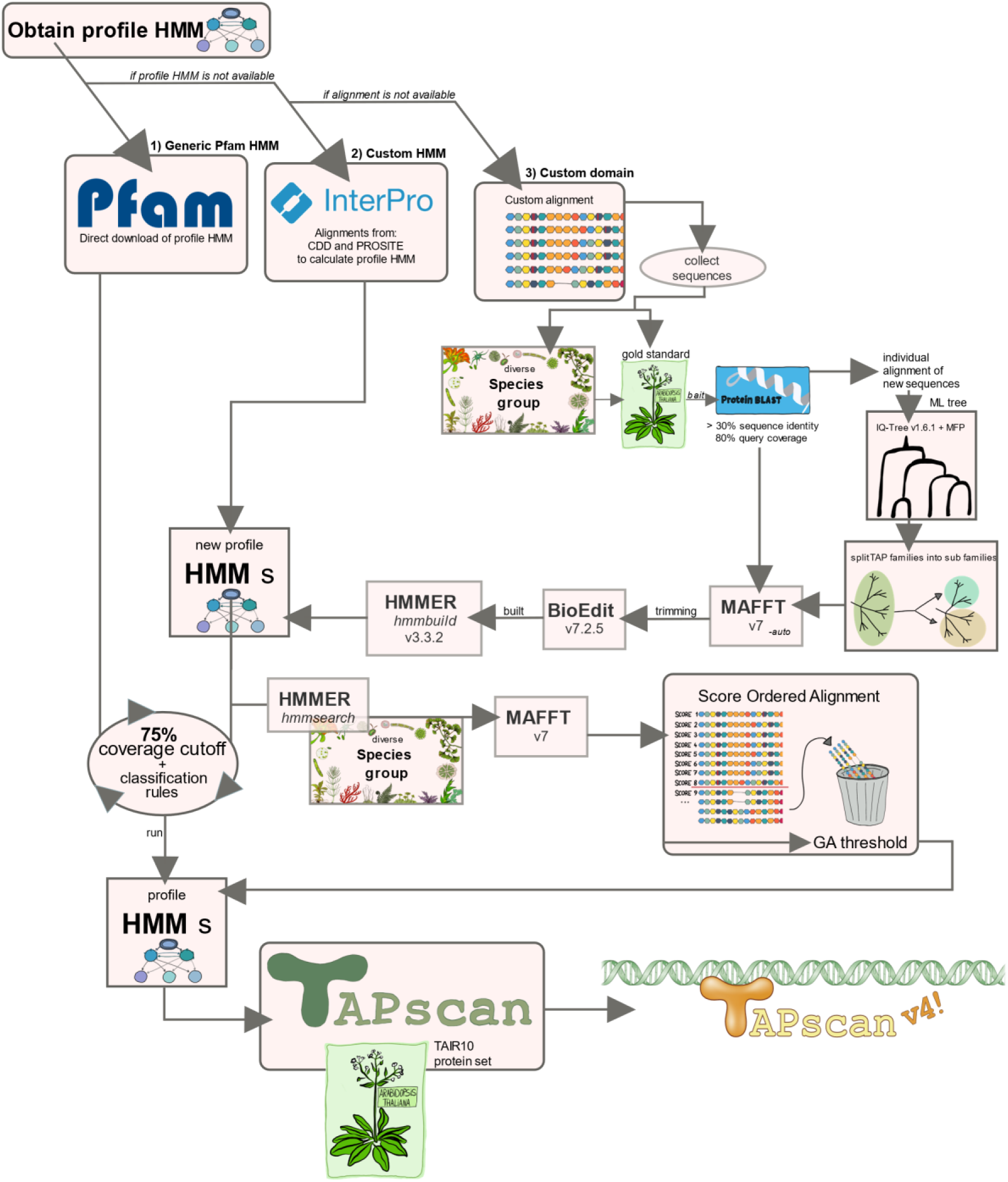
Methodology flow chart. Flow chart representing the process of adding new TAP (sub)families to TAPscan. If an entry was found within the Pfam or another database included in InterPro (= custom HMM), the profile was downloaded, manually edited if needed and then included into TAPscan. In case no entry was found, a custom alignment + HMM (= custom domain) was generated for the respective domain and included into TAPscan.

#### 2.2.1 Obtaining a profile Hidden Markov Model

To acquire a new profile HMM to detect a newly added TAP family, three sources were considered. First, searches using InterProScan 86.0 were performed using protein sequences belonging to the TAP family to be added (Blum et al., 2021). As first choice, if the profile was present in the Pfam database (version 34.0, March 2021), the profile was directly downloaded from the website (generic Pfam profile) (Mistry et al., 2021). As second choice, if an entry of the conserved domains database (CDD) or the PROSITE database was detected, the corresponding alignment was downloaded and used to calculate a custom HMM (S. Lu et al., 2020; Sigrist et al., 2012). If none of the searches matched any domain, a custom domain was created by first generating a custom alignment and then translating this into a custom HMM. Hence, HMMs used for TAPscan either are represented by generic Pfam profiles, are custom HMMs based on a published alignment, or are custom domains (alignment + HMM).

Sequences for custom alignments were collected either by using the sequences provided in the gold standard publication, by using sequences from DS1 (Table S1) or by additionally performing BLASTp searches with selected “bait” sequences listed in the gold standard to collect further appropriate sequences (Altschul, 1997). Using BLASTp, detected sequences were required to possess at least 30% sequence identity and 80% query coverage in order to avoid the twilight zone of protein alignments (Lang et al., 2010; Rost, 1999). If a subfamily of an existing TAP family was to be added, the previous profile HMM was adapted so that it only contained the sequence motif to detect the subfamily. The information about the origin of each newly added domain profile HMM can be found in Table S3 (see File S1 for all profile HMMs and File S2 for custom alignments).

The downloaded sequences or alignments (i.e. custom alignments and alignments from the CDD or PROSITE database) were used to construct multiple sequence alignments using MAFFT version 7 with the --auto option (Katoh et al., 2019). BioEdit v7.2.5 was used to manually trim the alignments to exclusively represent the domain motif in question (Hall, 1999). To calculate the final profile HMMs, hmmbuild provided in the HMMER software package was used (Eddy, 1998). All tools were run on servers which use Ubuntu 20.04.3 LTS.

#### 2.2.2 Definition of gathering thresholds

Each sequence to be annotated is scanned for the presence of specific protein domains during a TAPscan run, and the significance of these matches is indicated using the “sequence bit score” provided by hmmsearch. This score is the log-odds score for each sequence and is positively correlated with the significance of the detected hit (Eddy, 1998). The gathering-threshold (GA-threshold) is part of every profile HMM in TAPscan and specifies a reference bit score for the respective domain, and discards sequences that exhibit scores lower than this score. For each newly added custom profile HMM a suitable GA-threshold needs to be defined by the use of so-called score-ordered alignments (Wilhelmsson et al., 2017). In these alignments, sequences are ordered descending according the sequence bit score. The ordered alignments are manually assessed, and the domain motifs and sequence similarities of the sequences compared. The final GA-threshold is set at the position where the sequence motif of a sequence deviates from the above, better-scoring sequences. Thus, the GA threshold for a given domain was defined based on the sequence bit score of the first sequence with a motif deviating from the top-scoring motifs. Furthermore, the corresponding “acc” value for every sequence is considered. The “acc” value is the “average posterior probability of the aligned target sequence residues”, indicating the expected accuracy for every position in the alignment (Eddy, 1998). The species group used to calculate the score-ordered alignments is described in section 2.1; the tool used to calculate the assemblies was MAFFT (Katoh et al., 2019). The final step was to perform TAPscan runs using *A. thaliana* (*E. siliculosus* in case of HD_TALE) with the included new profile HMMs using the newly defined GA-thresholds to test the updated complete TAPscan family set in terms of whether it would correctly detect all TAP family members. The final collection of profile HMMs can be found in File S1 and are included in the TAPscan v4 GitHub repository (https://github.com/Rensing-Lab/TAPscan-v4-website).

#### 2.2.3 Definition of coverage cutoffs and specific rules

As investigated in (Wilhelmsson et al., 2017) and confirmed here, 75% is a good general coverage cutoff value to set for new custom domains. This cutoff value refers to the percentage to which a sequence to be annotated must match a given profile HMM. Sequences that match less than this percentage are discarded, ensuring higher specificity in assignment. In addition, domain-specific classification rules were defined for each TAP family, i.e. the domains that should be present or absent in a protein in order to be assigned to the respective TAP family (File S3). For certain TAP families it was also necessary to apply changes to the TAPscan script. For instance, there are TAP families where the hits of two different subfamilies are combined (e.g., in the case of the family bZIP, which comprises the subfamilies bZIP1, bZIP2, bZIPCDD and bZIPAUREO; all hits of these subfamilies are merged and, in the output, referred to as the family bZIP). Such special rules are defined for the families bZIP, GARP_ARR-B and ET (cf. Figure 2). The final coverage cutoffs and classification rules can be found in File S4 and S3, respectively.

**Figure 2.**
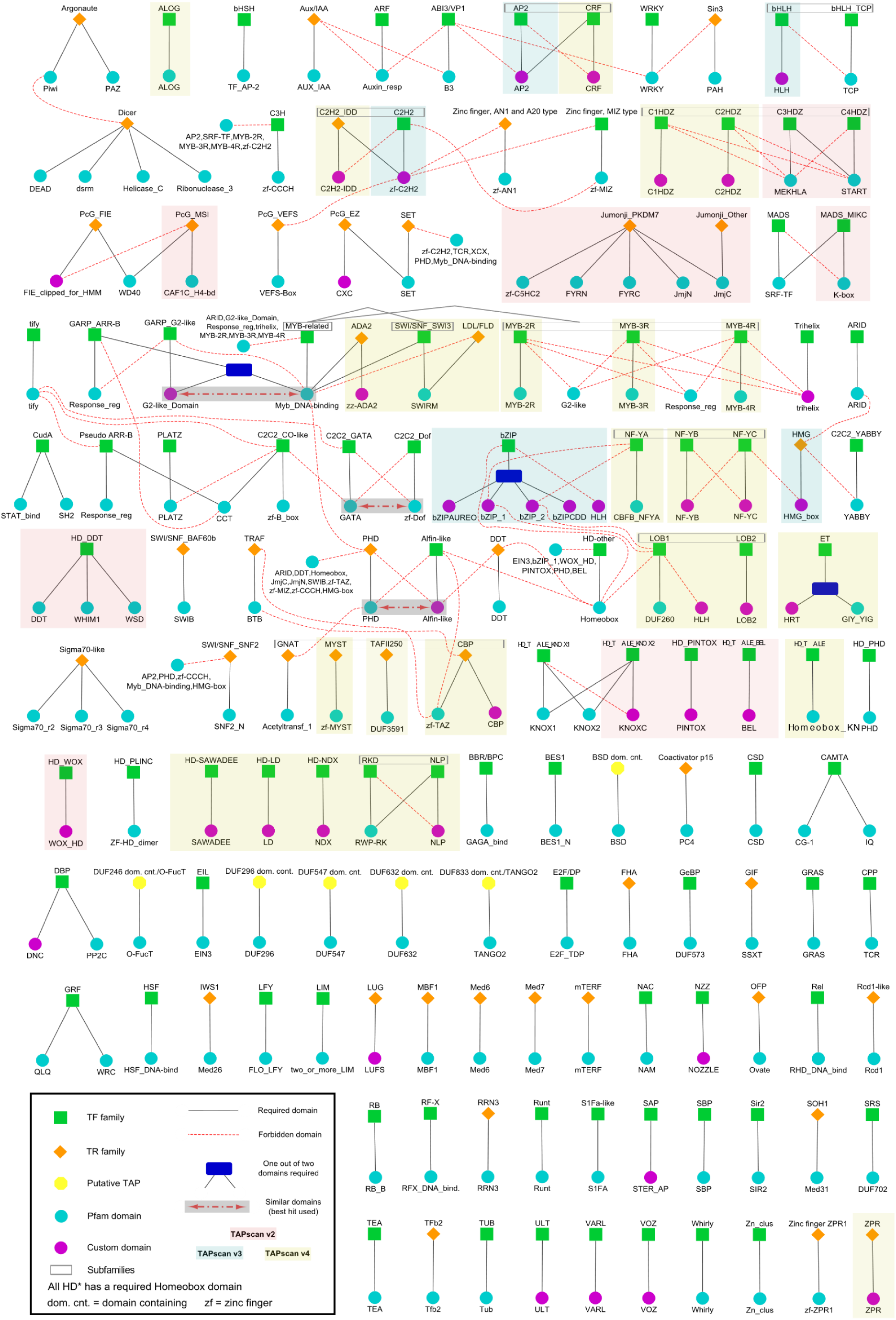
Overview of the TAPscan v4 rule set. An overview about the 138 TAP families with their corresponding classification rules is given, indicating which protein domains should or should not be present in order to assign a protein to a TAP family. The colors of squares, circles and diamonds indicate the type of TAP (TF, TR, or PT) and the origin of the domains (Pfam domain or custom HMM/domain). Custom HMMs (built from an Interpro alignment) and custom domains (based on a custom alignment; cf. Methods 2.1) are both summarized as “custom” here. Black lines indicate required (“should”) domains to define the TAP family, red lines indicate forbidden (“should not”) domains (that must not occur). A blue box specifies that one out of two domains must be detected, while a red dashed arrow in a gray box indicates that the better scoring hit of these two domains is used to assign the TAP family. TAP families highlighted by yellow background were added as part of the TAPscan update to v4. For the families highlighted in blue, changes were made as part of the update to TAPscan v3, while families highlighted in red were updated as part of the update to version 2. Grey boxes surrounding TAP families indicate subfamilies.

### 2.3 Phylogenetic support to define subfamilies

To ensure accurate separation between the newly defined subfamilies of the TAP families LBD and HDZ, phylogenetic trees were calculated (File S5). In both cases, DS1 (cf. Table S1) was used. First, TAPscan runs were performed using the previous TAPscan v3, i.e., with the TAP families not yet separated into subfamilies. The detected sequences of the former families ”AS2/LOB” and ”HD-Zip_I_II” were received using seqtk (https://github.com/lh3/seqtk). Since almost 900 sequences were detected for AS2/LOB using the complete species group, and to avoid a huge phylogenetic tree, sequences were deleted randomly so that 465 sequences were finally present. Multiple sequence alignments were calculated using MAFFT with the --auto option for both, AS2/LOB and HD-ZIP_I_II sequences (Katoh et al., 2019). BioEdit (Hall, 1999) was used to trim the alignments according to the respective gold standard and to delete intervals between the conserved blocks. The IQ-Tree multicore version 1.6.1 was used with the MFP (ModelFinder Plus) option to calculate maximum likelihood (ML) trees and to determine the best evolutionary models (Nguyen et al., 2015). For the AS2/LOB family the model JTT+R5 performed best and was chosen. For the HDZ tree, the best-performing model JTT+I+G4 was used. The trees (File S5) were visualized using FigTree v1.4.4 (https://github.com/rambaut/figtree). Based on the distribution in the trees, the included sequences were then divided into LOB1 and LOB2, and C1HDZ and C2HDZ, respectively, and used to build the profile HMMs of these subfamilies.

### 2.4 The TAPscan script

When executing TAPscan, after running *hmmsearch* the final annotation of the TAP families is performed by executing the TAPscan Perl script (available from https://github.com/Rensing-Lab/TAPscan-classify) to apply the defined classification rules and coverage values. After execution, three different outputs are generated. Output 1 indicates which domains were assigned to each sequence, and which TAP family this sequence was finally assigned to. The new additional output 3 follows the same structure as output 1 but additionally enables a differentiation of subfamilies. Output 2 provides a summary of how many sequences were assigned to each of the 138 TAP families.

During the update, the implementation of subfamilies was achieved by defining the new output 3, but also by adding specific rules to the TAPscan script to determine which TAP families should be divided into subfamilies. TAP families that contain subfamilies in TAPscan v4 are C2H2, bHLH, NF-Y, HDZ, HAT, LBD, MYB, MYB-related, RWP-RK and AP2 (Figure 2).

Furthermore, to detect the new TAP family ET using the two domains HRT and GIY_YIG, an additional rule was added to the script to assign the results of both domains to the ET family. Also, the TAP family CCAAT_Dr1 is no longer included in TAPscan v4, based on the recent classification presented by (Zanetti et al., 2017). Therefore, a specific rule in the script that prefers the CCAAT_Dr1 domain was deleted, as well as an obsolete rule for the former TAP family HDZIP.

### 2.5 Statistical tests, PCA and asymmetric Wagner parsimony

To compare the total numbers of TAPs in different phyla and to check whether certain TAP families are significantly more abundant in embryophytes, streptophyte algae or in other algae (Chlorophyta, Prasinodermophyta, Rhodophyta, Glaucophyta and Cryptophyta), statistical tests were performed. For this, Shapiro-Wilk normality tests were performed with R version 4.3.1 (R Core Team, 2022) to check for normal distribution (p-value ≤ 0.05). Then, according to the result, two-sample t-tests (for normally distributed data) or Wilcoxon rank-sum tests were performed using R (p-value ≤ 0.05) (Table S4). The resulting p-values were corrected for multiple testing using the Benjamini-Hochberg correction (Benjamini & Hochberg, 1995). Principal component analysis (PCA) was performed to examine the distribution of species based on their TAP complements (Figure 6). For the calculation of the PCA, R and the package ggplot2 were used (R Core Team, 2022; Wickham, 2016) based on the reduced DS2 described in section 2.1.

To calculate lineage-specific gains, losses, contractions and expansions of TAP families, the asymmetric Wagner parsimony with default settings as implemented in the count package was used (Csuos, 2010). The species tree (astral-33-new-renamed.tre) from (Leebens-Mack et al., 2019), calculated based on 410 single-copy genes, was modified to include only species that were investigated here (File S6). The reduced DS2 was used as described in section 2.1. For species being not present in the species tree, the closest relative was chosen. The results of this analysis are presented in Table S5.

### 2.6 TAPscan online resource and accessibility

All TAP data is available from the TAPscan v4 web application (https://tapscan.plantcode.cup.uni-freiburg.de). TAPs can be interactively explored either by family or by species, and sequences can be viewed and downloaded. TAPscan v4 is written in PHP, using the Laravel 8 Framework with a MySQL 8.0 database. It utilizes a containerized setup based on Docker. A Docker file is utilized to precisely configure the development environment for TAPscan v4, with finely-tuned integration with the Laravel Sail framework. Essential dependencies, including PHP 8.0, Composer, Node.js, and MySQL, are incorporated in the Docker file based on the Ubuntu 20.04 image. The webpage was set up with the usage of PHP 8.0, JavaScript, HTML with Bootstrap, and CSS3 for the development of dynamic web interface. The tree visualizations were generated using ETE toolkit (Huerta-Cepas et al., 2016). The code for the TAPscan v4 web application, and all data contained within, is also freely available under a GPLv3 license (https://github.com/Rensing-Lab/TAPscan-v4-website) for any users interested in running their own TAPscan application.

### 2.7 Pre-computed phylogenetic trees on the website

To aid the species view on the website, phylogenetic trees were calculated for each TAP family, a Neighbor-Joining (NJ) and/or a Maximum Likelihood (ML) tree. A total of 145 species belonging to the Archaeplastida were used, with a focus on embryophytes (Table S1, tab 3 “DS3 Genome Zoo”, column AG). In case of TAP families for which less than 1,000 hits were detected within these species, we proceeded similarly to section 2.3. The alignments were calculated using MAFFT using the -auto option and trimAl v1.4.rev22 was used to ensure consistent trimming, using a gap threshold of 50% (option -gt 0.5) and a similarity score lower than 0.001 (option -st 0.001) (Capella-Gutiérrez et al., 2009; Katoh et al., 2019). Quicktree v2.5 was used to calculate the NJ trees using 100 bootstrap replicates (Howe et al., 2002). IQ-Tree v1 was used for calculating the ML trees, using the ultrafast bootstrap option with 1,000 replicates, the parameter -m MFP to determine the best-fit substitution model and the parameter -alrt to perform SH-like approximate likelihood ratio tests also with 1,000 bootstrap replicates (Kalyaanamoorthy et al., 2017; Nguyen et al., 2015). If more than 1,000 hits were detected for a given TAP family, a procedure was applied to reduce the number of branches in the tree for brevity. For this purpose, the EMBOSS version 6.5.7 pairwise alignment application needleall was applied using a gap open penalty of 20.0 and gap extension of 0.2 (Rice et al., 2000). By applying needleall, pairwise percentage identities were calculated for each sequence comparison (Rice et al., 2000). For all pairwise sequences with percent identities above 90% one sequence was deleted, leading to a smaller data set. For these datasets, alignments and trees were calculated as outlined above. The tree visualizations were generated using ETE toolkit (Huerta-Cepas et al., 2016).

## 3. Results and Discussion

### 3.1 TAPscan v4 comprises 18 additional (sub)families

TAPscan is a comprehensive tool for annotation of TAPs and thus ideally applicable for analyses of total TAP complements, specific TAP families, and their lineage-specific evolution. With version 2 and 3, the number of included TAP families as well as the sensitivity and specificity in the detection of TAP families by TAPscan had been improved as compared to v1 (Petroll et al., 2021; Wilhelmsson et al., 2017). The first version of TAPscan was published in 2010 (Lang et al., 2010), the second, expanded version, TAPscan v2 was published seven years later (Wilhelmsson et al., 2017). In 2021, TAPscan v3 with increased sensitivity regarding the detection of TAPs in algae was published (Petroll et al., 2021). TAPscan has been cited several hundred times, with citations referring either to results of the performed phylogenetic and comparative analyses of TAP complements, or referring to analyses where the provided annotated TAP datasets were applied to analyses like gene regulation and protein domain analyses, to functional analyses of TAP families, especially in relation to the evolution of morphological complexity, as well as to phylogenetic studies encompassing aspects of land plant and algal evolution. It should be noted that the TAPscan methodology has been adapted to other questions, for example in a domain analysis of the 1,000 thousand plant transcriptomes project (Leebens-Mack et al., 2019), and has also been reverse engineered for several studies, for example (Wang et al., 2019).

For TAPscan v4, the goal was to enable a more comprehensive detection of several TAP families with special focus on adding and editing subfamilies. The aim was to improve the overall sensitivity and specificity of TAPscan. In total, the update to version 4 encompasses the addition of 18 new TAP (sub)families, the update and renaming of selected (sub)families, and the deletion of two families. Of note, all new families are detected with 100% sensitivity and specificity as compared to the respective gold standard (Table S3 and Table S6). In total, TAPscan v4 enables the annotation of 138 different TAP families using 155 domains (Figure 2).

The domains of five of the 18 new TAP (sub)families were received from the Pfam database, namely for ALOG, MYST, HD_TALE, TAF_II_250, and ET (Mistry et al., 2021). TAF_II_250 and MYST are among at least four histone acetyltransferases found in eukaryotes, of which all four are now included in TAPscan v4 (Uhrig et al., 2017). In previous TAPscan versions, the TF family HRT (*Hordeum* repressor of transcription) was included based on a publication by (Raventós et al., 1998). After reviewing the more recent publication by (Tedeschi et al., 2019), an additional domain for the detection of this TF family was added and the family was renamed ET (EFFECTORS OF TRANSCRIPTION). The previous HRT family was detected with a sensitivity of 67% in *A. thaliana,* which now was increased to a sensitivity of 100% using TAPscan v4 (Table S6).

Heterodimerizing TALE homeodomain (HD) TFs are found in Metazoa as well as Archaeplastida and have been suggested to be an ancestral eukaryotic mechanism for controlling the haploid-to-diploid life cycle transition, and to have diversified in lineages with complex multicellularity (Dierschke et al., 2021; Joo et al., 2018). In the Chloroplastida, the KNOTTED-like (KNOX) homeodomain and the BEL1-like (BEL) homeodomain function as heterodimers (Hamant & Pautot, 2010; Joo et al., 2018). Interestingly, TALE TFs are also present in brown algae (Lee et al., 2008), with the two TALE_HDs SAMSARA and OUROBOROS heterodimerizing and thus controlling life cycle transitions (Arun et al., 2019). However, the full HD TALE complement of brown algae could not be detected with TAPscan V3. Moreover, divergent (sub)family classification led to conflicting evolutionary interpretation in the literature (cf. Table S5, tab 1, lines 62+68-79/columns G-AG). To enable the accurate classification of all TALE HDs with TAPscan v4, also those with deviating domain motives compared to the Chloroplastida, the HD_TALE subfamily was included by adding an additional domain (Homeobox_KN from Pfam). All three TALE HDs of *E. siliculosus* can be classified by TAPscan v4 as HD_TALE with increased sensitivity and specificity of 100%. For clarity, the already existing subfamilies have been renamed HD_TALE_KNOX and HD_TALE_BEL. The HD TF superfamily is a so-called pan-eukaryotic family (Catarino et al., 2016). The superfamily can be divided into 11 subfamilies (using HD_TALE_KNOX and HD_TALE_BEL as two independent subfamilies of HD_TALE, and HDZ as another subfamily), of which eight were already present in previous TAPscan versions (Mukherjee et al., 2009). To enable the detection of all 11 subfamilies with TAPscan v4, the HD-LD, HD-NDX and HD-SAWADEE subfamilies were included. Furthermore, the ZPR family which resulted from a duplication of a C3HDZ paralog was also added to the TAPscan family set (Floyd et al., 2014).

Three additional TAP families were added through alignments provided by the Prosite and CDD databases (custom HMMs): CBP, ADA2, and NLP. CBP is a histone acetyltransferase and ADA2 is one of three subfamilies belonging to the SWIRM domain proteins (Gao et al., 2012; Uhrig et al., 2017). Based on (Chardin et al., 2014), the RWP-RK TF family, which was already integrated in previous TAPscan versions, was divided into the two subfamilies RKD and NLP. To enable this differentiation in TAPscan, the NLP subfamily was added and the RWP-RK family was renamed RKD.

For eight of the 18 new (sub)families, no existing entry was found in any of the databases. Therefore, custom domains, i.e., domains based on a manually curated alignment, were calculated (cf. Methods). These are C2H2-IDD, CRF, ZPR, HD-LD, HD-NDX, HD-SAWADEE, the LBD family which was split into the subfamilies LOB1 and LOB2, and the HDZ family for which the subfamilies C1HDZ and C2HDZ were defined. In case of the HDZ family, where family HD-ZIP_I_II was divided into the subfamilies C1HDZ and C2HDZ, the sensitivity in detecting the C1HDZ subfamily was improved from 57% using previous TAPscan versions to 100% in v4. Based on two publications from 2020, the AS2/LOB family was renamed LBD and divided into the two sub families LOB1 and LOB2, with a custom domain created for LOB2 to allow this differentiation (Huang et al., 2020; Zhang et al., 2020). To subdivide AS2/LOB und HD-ZIP_I_II to LOB1, LOB2, C1HDZ and C2HDZ, respectively, phylogenetic trees based on 265 sequences for AS2/LOB and 334 sequences for HD-ZIP_I_II were calculated (File S5). Furthermore, C2H2-IDD and CRF were added as subfamilies for the TF families C2H2 and AP2, respectively, to ensure higher accuracy in the detection of these families.

Finally, the families MYB, LDL/FLD and NF-Y were updated. In these cases, no additional domains were added, but the classification rules were adjusted. MYB can be divided into the three subfamilies MYB-2R, MYB-3R, and MYB-4R based on the number of imperfect sequence repeats of the MYB domain (Cao et al., 2020). The classification rules of the original SWI/SNF_SWI3 family from previous TAPscan versions were updated based on current literature, so that the classification is now based on revised rules and SWI/SNF_SWI3 is now a subfamily of the MYB-related family (Gao et al., 2012; Genau et al., 2021). In addition, using the original rules to detect the SWI/SNF_SWI3 family, the LDL/FLD TR family can now be detected with TAPscan v4. The TF families belonging to the CCAAT group of previous TAPscan versions have been renamed to NF-YA, NF-YB and NF-YC and the CCAAT_Dr1 subfamily has been deleted based on a current gold standard publication (Zanetti et al., 2017). Furthermore, the domain zf-TAZ used in previous versions for the detection of the TAZ family is now used for the detection of the CBP subfamily based on the classifications of (Uhrig et al., 2017; Yuan & Giordano, 2002). To account for subfamily classification, an additional TAPscan output file is now generated. As an example, the annotated TAP (sub)families and sequence IDs for *A. thaliana* in form of the original output format (output 1) and the new additional file format (output 3), as well as a summary of how many hits were detected per TAP family (output 2) can be found in Table S6 together with a comparison of sensitivity and specificity values.

### 3.2 TAPscan v4: enhanced webpage functionality

As part of the update to TAPscan v4, not only the complete set of TAP families, but also the associated website and accessibility, have been revised and improved. The updated v4 script is also made available (under a GPLv3 license) to enable the use of TAPscan independently as a comprehensive tool on user-provided data (https://github.com/Rensing-Lab/TAPscan-classify). Furthermore, the TAPscan classification script has also been made available as a Galaxy tool (https://toolshed.g2.bx.psu.edu/), and is freely available to use from the European Galaxy server (https://usegalaxy.eu).

The updated version of the TAPscan web interface makes the annotated TAP data of all 678 species from our Genome Zoo available (https://tapscan.plantcode.cup.uni-freiburg.de). The TAPscan v4 website provides both a species view and a family view (Figure 3). The family view provides detailed information on all 138 different TAP families. Subfamilies are also included on this page, and are prefixed with the family name. In addition, at least one phylogenetic tree has been calculated for each family, a ML and/or a NJ tree, which are interactively displayed directly on the website, and can be downloaded in Newick format. The species view is a searchable taxonomic tree structure of all species that is based on NCBI taxonomy (Schoch et al., 2020). By selecting specific species, the TAP data for that species can be viewed and downloaded, and links are provided to search for each protein on PLAZA (Van Bel et al., 2022). A general search function was implemented to search for specific sequence IDs, species names and TAP families.

**Figure 3.**
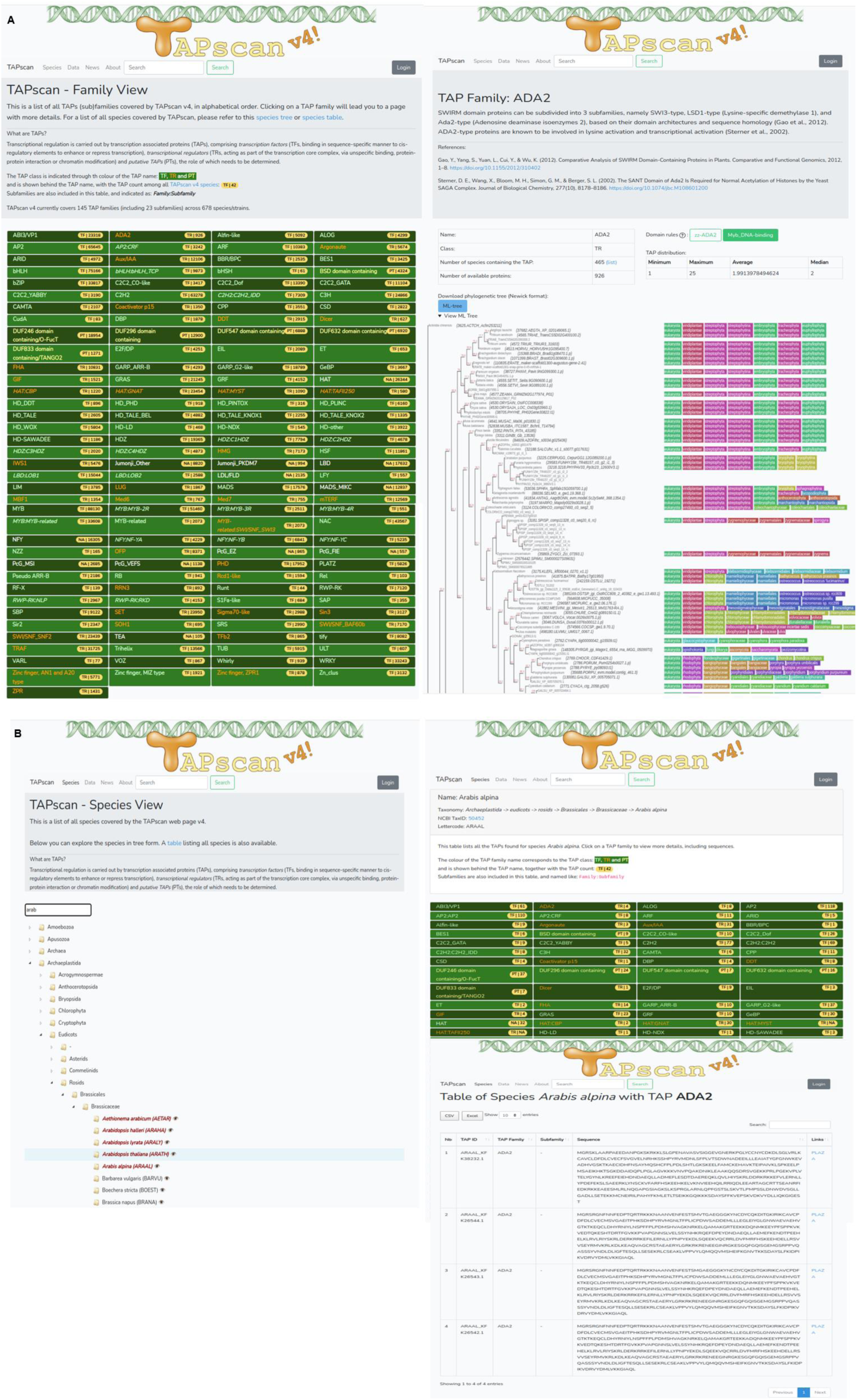
TAPscan v4 website. **A)** The TAPscan website includes a list of TAP families on its homepage (family view, left), and a detailed family page featuring family information and domain rules, and including a phylogenetic tree (right) that is decorated with NCBI taxonomy information, Subfamilies are indicated as: family:subfamily. **B)** The species view page includes an interactive and searchable species tree (left), leading to individual species pages showing a table of all detected TAPs (top right) which in turn leads to pages where individual TAP information can be viewed and downloaded (bottom right).

Data management has also been improved as data can now be uploaded and modified directly via an admin dashboard in the web interface. Data that must be provided includes the subfamily classification output from the TAPscan script, a list of species names and protein fasta files, as well as the rule set which was used for the classification. Detailed documentation about installing TAPscan v4 and provisioning data for inclusion is available on GitHub (https://github.com/Rensing-Lab/TAPscan-v4-website)

The availability of the stand-alone classification script (either as a command-line or Galaxy tool), enables use of the tool on novel data sets of interest, and to use it beyond TAP classification, by adding/replacing HMMs and the rule definition file. The TAPscan domain-based classification scheme indeed has already been applied in the past to other projects, for example the 1,000 transcriptomes project (Leebens-Mack et al., 2019) or the analyses of novel Zygnematophyceae genomes (Cheng et al., 2019).

This TAPscan version also for the first time features the release of the underlying “Genome Zoo” dataset (cf. Methods and Table S1, https://github.com/Rensing-Lab/Genome-Zoo). This dataset draws on published genomic datasets and represents non-redundant protein sets for a range of nearly 700 species. It is focused on Archaeplastida, comprising 70% of the species, and here again with a focus on non-seed plants and streptophyte algae (10% of the Archaeplastida species incorporated), that are the topic organisms of the priority program MAdLand (https://madland.science/). However, it also features a large range of phylogenetically diverse species from animals, fungi, SAR group, bacteria and archaea (Figure 4). They include model organisms such as yeast, *E. coli* or mouse, and cover a diverse set of lineages that may serve as outgroups in phylogenetic analyses that focus on Archaeplastida. The Genome Zoo data set (Table S1, DS3) provides non-redundant protein information (each species is represented by a genomic dataset, and each locus by its representative isoform) and allows for straight forward species information in phylogenetic trees via the 5-letter suffix species code (cf. Methods, Figures 3b and 6).

**Figure 4.**
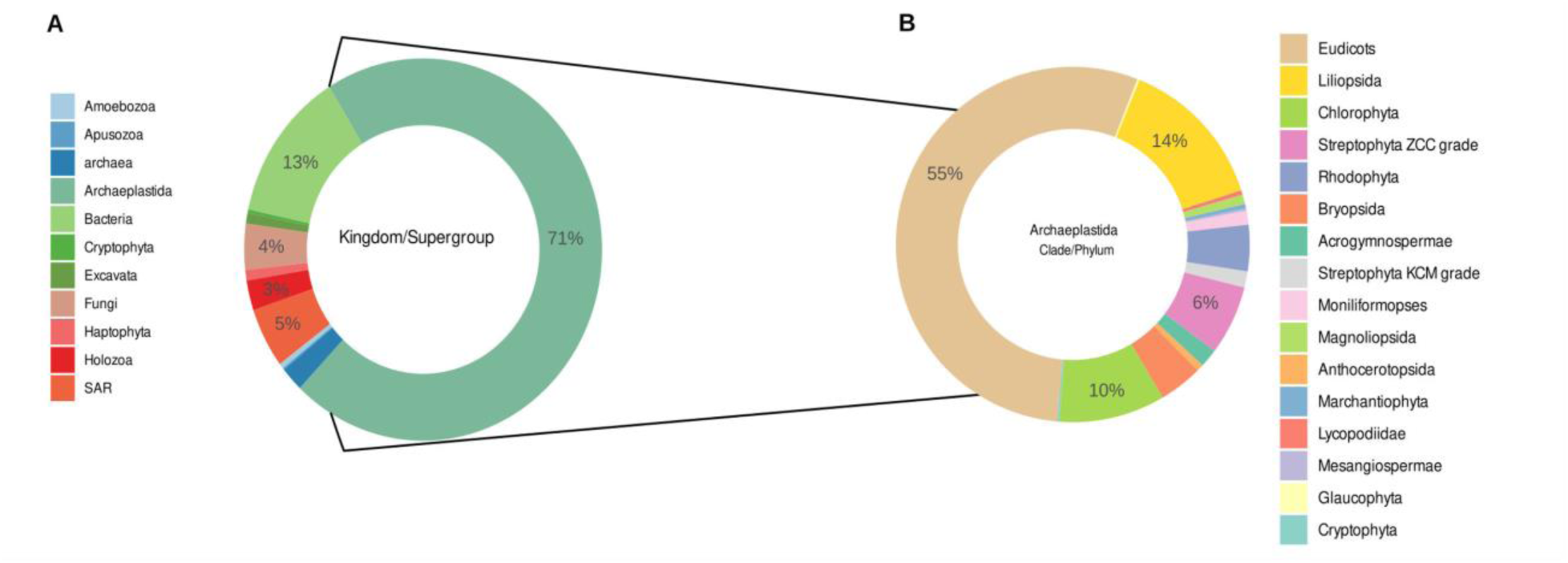
Species distribution in MAdLandDB/Genome Zoo. **A)** Distribution across kingdom/supergroup, **B)** Distribution across different phyla belonging to the Archaeplastida. The database has a focus on Archaeplastida, and within them on streptophyte algae and non-seed plants.

### 3.3 Gradual increase of TAP complement complexity during evolution of the green lineage

To investigate the evolution of TAP families in the Chloroplastida, TAPscan v4 was applied and members of 91 TF Families, 41 TR and 6 PT families in 37 species (Table S1, DS2) were identified. In general, most TAP families comprise significantly more members in embryophytes as compared to algae (Figure 5), indicating an increase of TAP family abundance during the evolution of the green lineage. Indeed, previous analyses have shown that a stepwise increase of TAPs, particularly TF family members, occurred during the evolution of Chloroplastida prior to the establishment of land plants (Catarino et al., 2016; Lang et al., 2010). It was also demonstrated that there is a general trend that the number of TAPs increases with the morphological complexity of an organism (Bowman et al., 2017; Lang et al., 2010). In (Lang et al., 2010), it was shown that the number of different cell types (used as a proxy for morphological complexity), is positively correlated with the total number of TAPs (and in particular TFs) encoded by Chloroplastida genomes. Morphological adaptations such as the occurrence of polyplastidy, a general increase in the number of cell types, the land plant-like cell wall, stomata (Harris et al., 2020), and vascular (K.-J. Lu et al., 2020) tissue have evolved during terrestrialization and enlarged the morphological complexity of plants (J. de Vries & Archibald, 2018; Jiao et al., 2020; Lang et al., 2010; Rensing, 2018). With the morphological traits, the underlying signaling pathways mediated via diversified TAP complements also diversified (J. de Vries & Archibald, 2018; Lang et al., 2010). Interestingly, the unicellular Zygnematophyceae species *Spirogloea muscicola* and *Penium margaritaceum* take an intermediate position and show, just like the embryophytes, increased amounts of a large part of TAP families (Figure 5). Most likely, similar to e.g. *Chara braunii*, they have secondarily increased their TAP complement (Cheng et al., 2019; Jiao et al., 2020; Nishiyama et al., 2018). At the same time, bryophytes such as the liverwort *Marchantia polymorpha* and the hornwort *Anthoceros agrestis* have apparently gone through secondary gene loss (F.-W. Li et al., 2020; Puttick et al., 2018, Harris et al., 2022). Gene family expansions and whole genome duplications (WGDs) were indispensable for the complexity evolution of land plants, since paralogs were retained, and numerous new functions and traits were established through sub- and neofunctionalization (Rensing, 2014, 2020).

**Figure 5.**
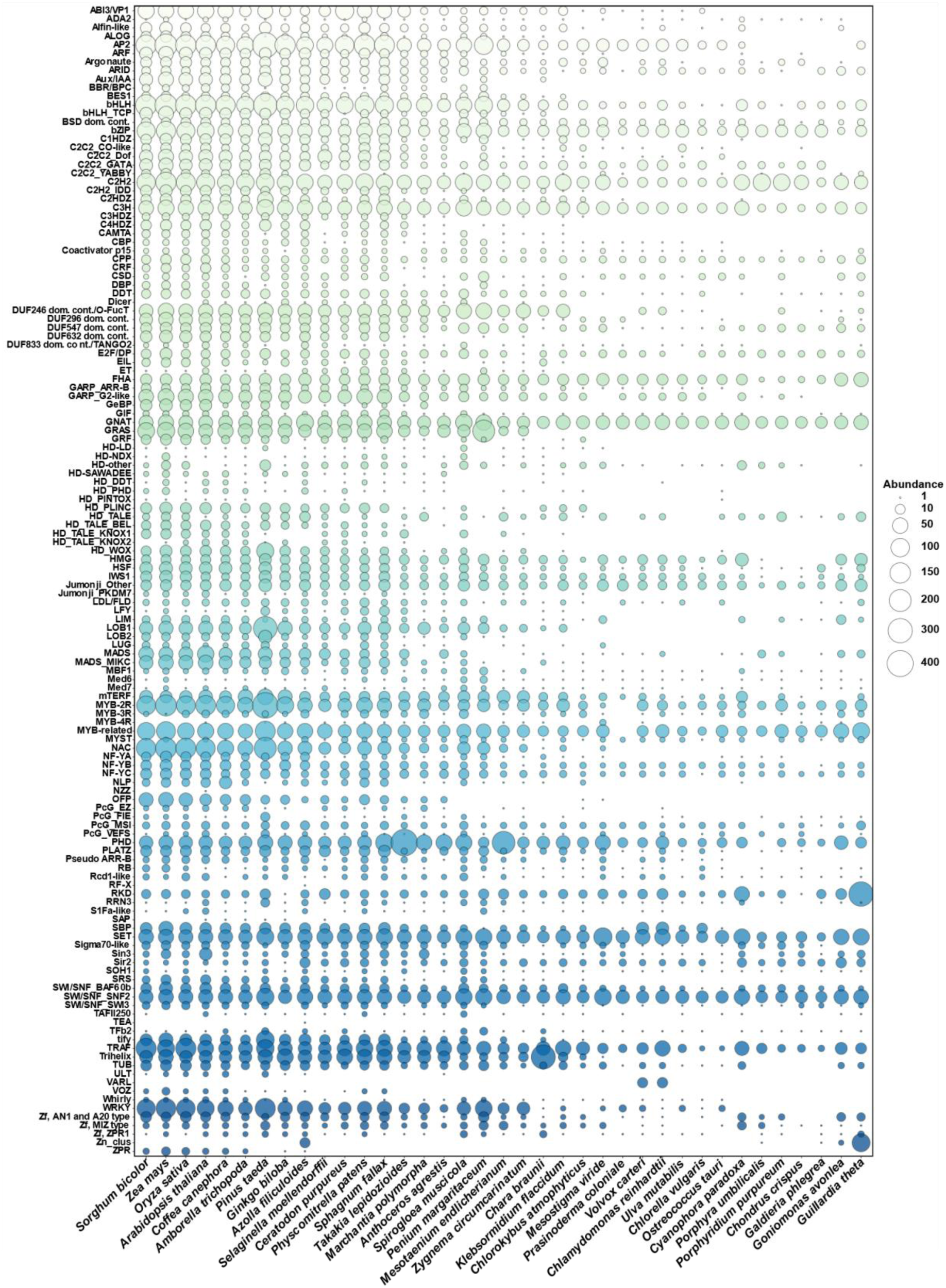
Bubble chart illustrating the distribution of TAP families occurring in the Chloroplastida. The numbers per each of 134 TAP families that occur in at least one of the species were plotted (size of the bubble corresponding to the number of family members). The species were ordered according to the species tree (cf. Methods).

### 3.4 High diversity of the TAP complements of streptophyte algae

To investigate the variance of TAP complements, a PCA was calculated (Figure 6) using 23 species and data from 134 TAP families (using only TAP families with at least one member across the investigated species). A general trend can be observed that the first PC divides species based on their total number of TAPs, with *Oryza sativa* containing 2,478 total TAPs and *Chlorella vulgaris* 213 TAPs on opposite sides. Overall, for the MRCAs of embryophytes and angiosperms, significant gains and expansions of TAP families can be inferred, which is also reflected by the comparatively high numbers of total TAPs in these species (Lang et al., 2010; Leebens-Mack et al., 2019). In addition, it can be observed that the species belonging to the Chlorophyta cluster, indicating similar TAP complements within this phylum.

**Figure 6.**
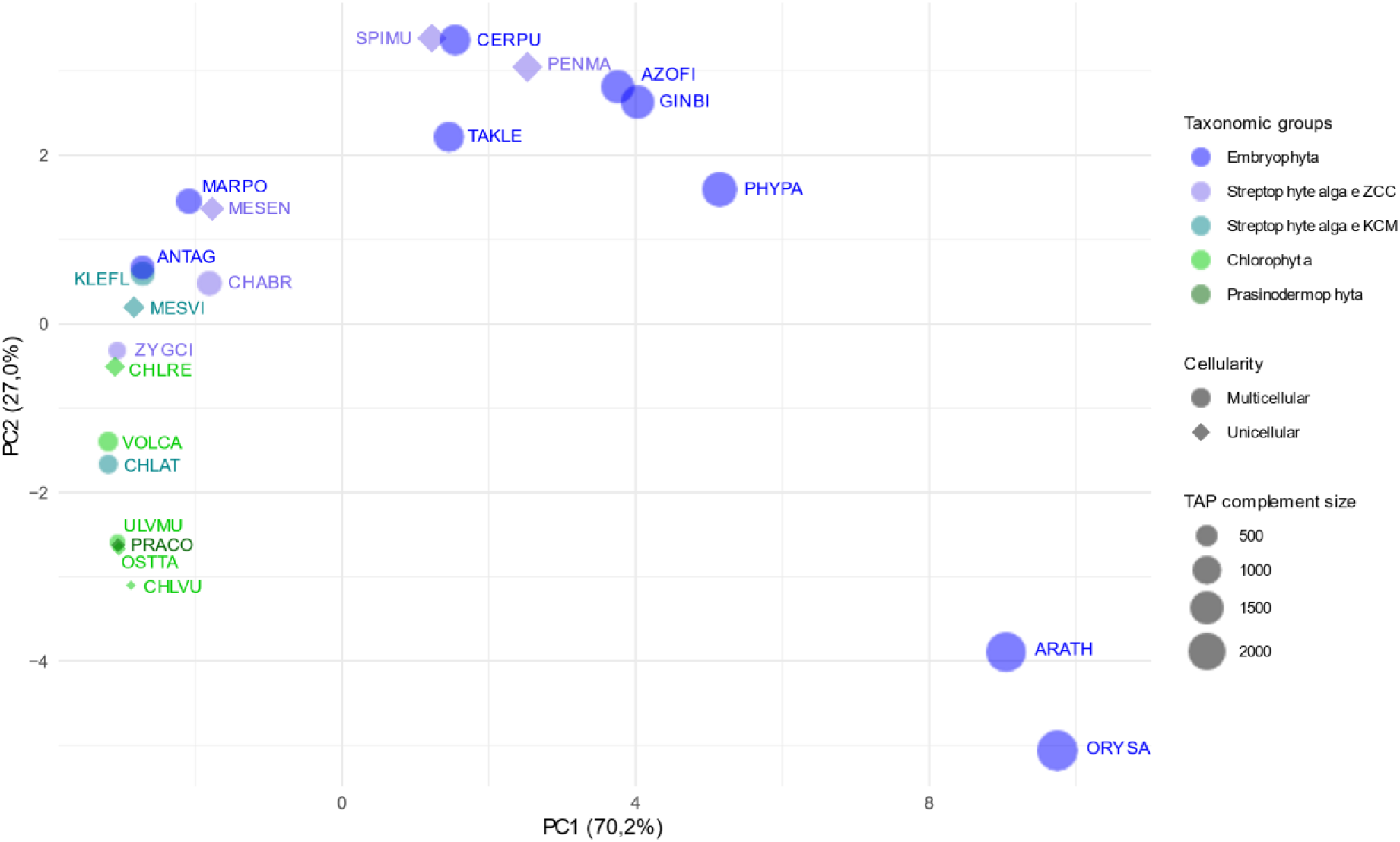
PCA analysis of the number of TAP families in the Chloroplastida with annotation of taxonomic groups. Data for a set of 134 TAP families of 23 species of Chloroplastida (Embryophyta, streptophyte algae ZCC and KCM grades, Chlorophyta and Prasinodermophyta). The x-axis shows the first principal component (PC), explaining 70.2% of the variance. The y-axis shows the second PC (27.0% of variance). Circles or diamonds indicate the cellularity of an organism, the size of the shapes indicate the total TAP complement size. Species names are abbreviated in the Genome Zoo five-letter code and are color-coded according to taxonomic groups (cf. Methods/Table S1).

In case of the streptophyte algae, species belonging to the KCM grade are clustering closely to the remaining algae, i.e., Chlorophyta and Prasinodermophyta, in contrast to species belonging to the ZCC grade which are spread across the PCA and in general cluster more closely with species belonging to the Embryophyta. Thus, the similar TAP complement is consistent with the branching order and with several unique characteristics that species belonging to the ZCC grade share with land plants, including more complex regulatory and developmental pathways (J. de Vries et al., 2016; Nishiyama et al., 2018; Rensing, 2020). Of note, two species (*Zygnema circumcarinatum* and *Mesotaenium endlicherianum* gene model V2, (Dadras et al., 2023)) of the Zygnematophyceae group cluster with bryophytes that did not feature WGDs, and with species belonging to the KCM grade, while *Penium margaritaceum* and *Sprirogloea muscicola* cluster with other species belonging to the Embryophyta (subject to WGD). Indeed, it has been remarked upon that *S. muscicola* was subject to a WGD event (Wang et al., 2019) (the only one so far detected in streptophyte algae), resulting in the triplication of its genetic material. This might have resulted in paralogs of certain TAP families and retention of these with subsequent sub- or neofunctionalization, and therefore a higher total number of TAPs.

*P. margaritaceum* features significant expansions of its entire genome (with a size of more than 4 Gbp), the number of protein-coding genes, and consequently also its TAP complement (Jiao et al., 2020). No WGD was detected in *P. margaritaceum*, but small-scale gene duplications of specific TAP families were observed (Jiao et al., 2020). One of these massive expansions was shown in (Jiao et al., 2020) for the GRAS TF family and can be confirmed as significant by comparing streptophyte algae with embryophytes (*p(fdr)* = 0.0152, Wilcoxon rank-sum test; Table S4). Based on various analyses (e.g. (Cheng et al., 2019; Grosche et al., 2018; Jiao et al., 2020)), it was suggested that the GRAS TF family evolved in the common ancestor of the Anydrophyta (Rensing, 2020), with an especially strong expansion in *P. margaritaceum*. In land plants, these proteins are involved in processes specific to complex multicellular organisms, in stress response and in the regulation of arbuscular mycorrhizal symbiosis (Cheng et al., 2019; Grosche et al., 2018; Jiao et al., 2020). The GRAS proteins independently expanded in streptophyte algae likely exhibit a functional divergence to those of the Embryophyta.

All 95 TAP families present in *P. margaritaceum* are also present in embryophytes. This suggests that the grouping of *P. margaritaceum* is largely due to the large number of total TAPs, resulting mainly from the expansion of GRAS TFs, while the detected TAP families are similar to those of embryophytes. In contrast, in *S. margaritaceum* there is no specific TAP family that is massively expanded. In *S. muscicola*, 100 different TAP families are present, all coincident with the TAP families found in embryophytes. TAPs represent a larger overall percentage of the proteome compared to *P. margaritaceum*, highlighting the expansion of multiple families as a result of the genome triplication event. This trend is confirmed by the performed analysis of evolutionary events using asymmetric Wagner parsimony: 19 species-specific expansions were detected in *S. muscicola* (Table S5). Interestingly, even with the small total TAP complement size and with grouping separately from the embryophytes in the PCA, *Z. circumcarinatum* exhibits the highest similarity within the streptophyte algae in terms of TAP families occurring in the Embryophyta, namely 104 TAP families. These findings indicate that a secondary reduction in morphological complexity does not generally correlate with a reduction of the TAP family complement, since *P. margaritaceum*, *S. muscicola* and *M. endlicherianum* are unicellular species and *Z. circumcarinatum* is multicellular (Feng et al., 2024; Jiao et al., 2020). This also aligns with the concept that the recurrent gain of multicellularity observed in Zygnematophyta — but also Klebsormidiophyceae — builds on an ancient genetic set that lingers also in unicellular species (Bierenbroodspot et al., 2024; Hess et al., 2022, Feng et al., 2024).

When observing the numbers of TAP families in *M. endlicherianum* and *C. braunii* an increased amount of specific TF families is striking, with 227 members (on average 42 members) of the TR PHD in *M. endlicherianum* and 287 members (on average 23 members) of Trihelix in *C. braunii*. The trihelix expansion was described in (Nishiyama et al., 2018) and these proteins were found to be expressed in all reproductive tissues, most of them in the oogonia and antheridia. Thus, a specific role in sexual reproduction, particularly in the antheridia, was suggested (Nishiyama et al., 2018). The bryophytes *M. polymorpha* and *A. agrestis* are grouping with species belonging to Chlorophyta, Prasinodermophyta, and streptophyte algae, rather than grouping with the Embryophyta, due to secondary reduction of their gene complement (Harris et al., 2022), respectively absence of WGD, as outlined above.

### 3.5 Significant gains in streptophyte algae and expansions in land plants

By comparing the total number of TAPs in embryophytes and streptophyte algae, significantly more TAPs (*p(fdr)* = 0.000361, t-test, Table S4) were detected in embryophytes than in streptophyte algae. When comparing further algae, namely Chlorophyta, Rhodophyta, Glaucophyta, Prasinodermophyta, and Cryptophyta, with streptophyte algae the same trend was observed (*p(fdr)* = 0.001848, Wilcoxon rank-sum test). 29 individual TAP families were identified as significantly expanded in the streptophyte algae as compared to other algae. 68 TAP families were detected as significantly increased in the embryophytes as compared to streptophyte algae. The TF family trihelix was found to be significantly increased in the streptophyte algae, as function of the massive expansion of this family in *C. braunii*. When comparing algae and embryophytes, 73 TAP families were found to be significantly increased in land plants. In (Leebens-Mack et al., 2019), the majority of TAP family expansions (18 in total) were detected during the transition from streptophyte algae to bryophytes. This conclusion is consistent with the statistical tests performed here, where significantly higher abundances were identified in embryophytes as compared to streptophyte algae, and in those as compared with other algae.

Based on the TAP complement analysis within the Streptophyta by (Bowman et al., 2017), not the general gain of TAP families was essential for terrestrialization, but rather expansions and diversifications of specific families. In particular, they mention in this context the families bHLH, NAC, GRAS, AP2, LBD and WRKY, which were argued to have been indispensable during the conquest of land (Bowman et al., 2017). In (Lang & Rensing, 2015), based on a PCA, TAP family size comparisons and a partial least square analysis, it was analyzed which TAP families show a correlation with the number of different cell types and therefore with the morphological complexity of an organism. It was shown that 28 different TAP families, on that basis, might be involved in the occurrence of multicellularity (Lang & Rensing, 2015). Among them are the families bHLH, C2H2, MADS and TRAF, which were also shown to be significantly expanded in embryophytes as compared to streptophyte algae in our analyses (Table S4). From the families mentioned by (Bowman et al., 2017) NAC was shown to be expanded in the Anydrophyta.

To further evaluate in which species or phyla expansions, contractions, gains or losses of specific TAP families have occurred, a species tree was constructed and asymmetric Wagner parsimony was applied to calculate these evolutionary events at each node of the tree (Csuos, 2010). For this, we used the species tree calculated by (Leebens-Mack et al., 2019) to ensure confident clustering, but scaled this tree down according to the species used here (Figure 7). The tree is outgroup-rooted using Cryptophyta as the sister lineage to the Archaeplastida (Adl et al., 2019; Strassert et al., 2021). Across all nodes, a total of 116 gains, 29 losses, 135 expansions, and 24 contractions of TAP families were detected. It must be kept in mind that the calculation and assignment of gains/losses and expansions/contractions are dependent on the sampling depth and thus some assignments will suffer from sampling bias. Most gains were observed within the streptophyte algae: 46 gains and therefore more than one third of all detected gains. Thus, the trend that most TAP-and especially TF-families were gained before the conquest of land (Bowman et al., 2017; Catarino et al., 2016; Wilhelmsson et al., 2017) was confirmed by our investigations.

**Figure 7.**
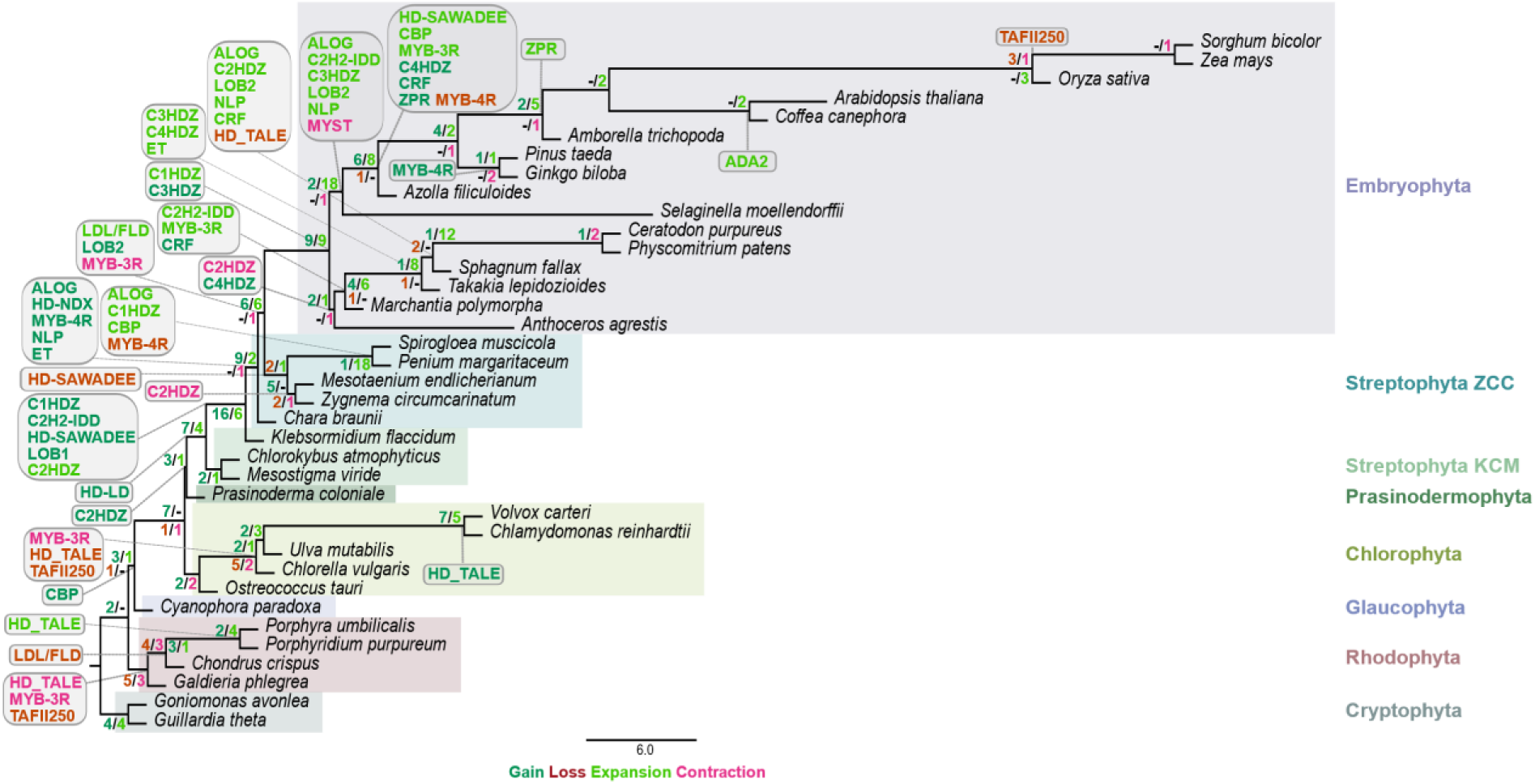
Species tree showing the total amounts of gains, losses, contractions, and expansions at each node. The tree is based on the species tree published by the one thousand plant transcriptomes (1kp) consortium (Leebens-Mack et al., 2019), is rooted using Cryptophyta as outgroup, and was scaled down to contain only the species used here. To calculate gains, losses, expansions and contractions, the asymmetric Wagner parsimony implemented in the count package was used (Csuos, 2010). The detected gains are shown in dark green, the expansions in light green, losses in dark red and contractions in pink. The highlighting indicates association with the phylum shown on the right in the same color. The rounded shapes show the detected events for the TAP families newly added or edited as part of the update to v4.

In contrast, most expansions occurred at the origin of the Embryophyta, i.e. during land plant evolution, a trend also shown by (Leebens-Mack et al., 2019). Expansions of 9 TAP families were detected in the MRCA of Embryophyta and another 18 in the MRCA of Tracheophyta. Interestingly, 20 expansions are detected within mosses. It should be noted that these might also be represent contractions in the other two bryophyte lineages, liverworts and hornworts. Yet, mosses are prone to WGD events, while the other two bryophyte lineages are not (Lang et al., 2018). *P. margaritaceum* and *S. muscicola* also show together a high number of 18 expansions - these massive expansions also evident by the clustering in the PCA (Figure 6) are probably due to massive small-scale duplications as well as WGD events.

### 3.6 Additional TAP families can be traced to an origin within streptophyte algae

A key aspect of the comprehensive update to TAPscan v4 was the definition of new subfamilies. These were of special interest in terms of defining their origins and in analyzing whether dividing TAP families into subfamilies allows more precise predictions about expansions and contractions. As part of the update, the TF family RWP-RK was divided into the two subfamilies RKD and NLP (Chardin et al., 2014; Wu et al., 2020). In the analyses conducted in (Wilhelmsson et al., 2017), at least one member of the RWP-RK family was detected in all but one species. However, no gain or expansion was detected for this family in the respective study (Wilhelmsson et al., 2017). Here, using the updated family classification, the RKD subfamily was detected in all species without any exception. The NLP subfamily, however, was shown to have been gained later, in the MRCA of the Phragmoplastophyta (cf. Table S5, tab 1, line 120). Therefore, the NLP subfamily was revealed to occur exclusively in the Phragmoplastophyta, and thus divergent and additional functions can be hypothesized compared to the RKD subfamily (Chardin et al., 2014). The phylogenetic analyses of (Sakuraba et al., 2022) shows that the NLP subfamily appears in terrestrial plant lineages and might have played a role in the adaption of nitrogen acquisition during the transition from water to land.

Similarly, in contrast to the TF family C2H2, for which members were detected in every species without exception, the C2H2-IDD subfamily was gained later within the MRCA of Klebsormidiales and Phragmoplastophyta and expanded subsequently in the Tracheophyta. (Prochetto & Reinheimer, 2020) were able to specify the origin of this subfamily to the common ancestor of the Streptophyta and specifically to the ancestor of the Klebsormidiophyceae and Phragmoplastophyta, consistent with the observed gain according to our analyses.

The MYB TF family was classified into MYB-2R, MYB-3R and MYB-4R to enable up-to-date classification in plants (Cao et al., 2020). In the previous analysis of (Wilhelmsson et al., 2017), the MYB TF family was present in every investigated species. Here, this distribution could be confirmed for MYB-2R and MYB-3R with few exceptions (potential losses). However, for the MYB-4R subfamily it can be shown that it was gained in the MRCA of Phragmoplastophyta and thus prior to terrestrialization. Interestingly, no members of MYB-4R were detected in *P. margaritaceum* and *S. muscicola*, therefore this subfamily was probably secondarily lost in these species, confirmed by the detected loss through asymmetric Wagner parsimony.

The TF family AS2/LOB was divided into the two subfamilies LOB1 and LOB2 based on recent literature (Zhang et al., 2020). In (Wilhelmsson et al., 2017) it was demonstrated that the AS2/LOB family was gained in the MRCA of Klebsormidiales and Phragmoplastophyta. For LOB1 this can be confirmed. LOB2, however, evolved in the MRCA of the Anydrophyta.

In (Romani et al., 2018) an analysis of the HDZ TF family in the Chloroplastida was performed. They suggest that the C2HDZ subfamily originated in chlorophytes and subsequently diverged into the C3HDZ and C4HDZ subfamilies and the C1HDZ subfamily. The assumption that C2HDZ is the ancestral subfamily can be confirmed by the results of the asymmetric Wagner parsimony performed here. However, the C2HDZ family was not detected in any species belonging to the Chlorophyta but in the prasinodermophyte *P. coloniale*. Hence, an origin in the MRCA of the Streptophyta was specified based on our parsimony results. It is also interesting that a gain of this subfamily was detected in *Cyanophora paradoxa*, belonging to the Glaucophytes. These findings suggest that the HDZ family possibly originated in the common ancestor of the Archaeplastida and was secondarily lost in several lineages. This suggestion would thus far predate the previously suspected origin and needs to be investigated further as further genomes become available.

As part of the update to TAPscan v4, the homeodomain TF families HD-LD, HD-NDX and HD-SAWADEE were included, so that all subfamilies belonging to the HD superfamily can be identified. Based on the analyses of (Mukherjee et al., 2009) and (Catarino et al., 2016) a correlation between the number and diversity of HD subfamilies and amounts of each subfamily with the complexity of an organism can be observed. In the analysis performed here, the family HD_TALE could be identified in all investigated phyla with the exception of a few species of Bryophyta, Prasinodermophyta and Chlorophyta. The newly added subfamily LD was identified as one of the ancestral subfamilies, in the MRCA of streptophytes. The PLINC and the SAWADEE subfamily were detected in the MRCA of Klebsormidiales and Phragmoplastophyta, as well as the subfamilies NDX and WOX. The TALE_BEL, DDT, TALE_KNOX_1, TALE_KNOX_2, PHD, and PINTOX subfamilies evolved afterwards, either in the MRCA of the Embryophyta or within the Embryophyta. Thus, the hypothesis that there is a coincidence between the number and diversity of HD members and the complexity of an organism is supported by the analyses conduced here. This trend has also been confirmed for the HD family during metazoan evolution, where the number and diversity of HD members increases with complexity evolution, while unicellular organisms possess comparatively smaller HD complements (Larroux et al., 2008; Sebé-Pedrós & de Mendoza, 2015). Expansion of the HD-TALE family coinciding with morphological complexity has also been found in brown algae recently (Denoeud et al., 2024). In summary, the progression of the emergence of the different subfamilies shows that before and concurrent with the conquest of land new TAP (sub)families evolved, probably enabling morphologically more complex organisms in adaptation to new environmental conditions and habitats. In (Lang et al., 2010), many TAPs previously classified as land plant-specific were shown to have been gained earlier, during the conquest of land. Subsequently, in (Wilhelmsson et al., 2017), a total of 26 TAP families were identified as gained in the MRCA of the Streptophyta, i.e., preceding terrestrialization. Here, by defining additional (sub)families in TAPscan v4 and with further genomes of streptophyte algae available, nine additional (sub)families were identified that were gained either in the MRCA of the Streptophyta or within streptophyte evolution. Thus, many TAPs that are important for life on land already evolved in streptophyte algae.

## 4. Conclusions

Here, we present the updated version of TAPscan v4 encompassing the addition of 18 new TAP (sub)families. We were able to add all new families with 100% sensitivity and specificity as compared to gold standards and improved overall accuracy by adding and defining new subfamilies. Combined with the public code release of TAPscan, the new version provides a framework for the community to perform comprehensive genome-wide annotation of TAPs or other protein families. Moreover, both TAPscan v4 and its underlying genomic dataset are now available via the galaxy framework and GitHub, free for academic use.

Previously it has been hypothesized that in the green lineage (similar to animals) increasing morphological complexity of an organism is correlated with increasing complexity of its TAP complement (Lang et al., 2010; Lang & Rensing, 2015). Based on a much higher number of species, this general trend indeed is observable. However, noteworthy exceptions include secondary gains and losses of TAPs in the Chlorophyta, streptophyte algae and bryophytes. Such deviations from the general trend are typically due to strong expansion of individual TAP families (like GRAS or Trihelix), or due to the presence/absence pattern of genome duplications.

Preceding the water-to-land transition, a gradual increase and thus progressive evolution of TAP families along the chain of ancestors in the streptophyte tree of life as well as an increase in diversity of many TAP families was previously inferred (Catarino et al., 2016; Lang et al., 2010; Wilhelmsson et al., 2017). Indeed, there are in general more TAPs in streptophyte algae than in other algae, and more in embryophytes again than in streptophyte algae. This trend was also confirmed in particular for the HD TF superfamily, with increasing morphological complexity the number and the amount of HD subfamilies increases. A diversified repertoire of TAPs in streptophyte algae suggests that the earliest land plants had a rich substrate on which evolution acted, sprouting new biological programs that were adaptive by responding to environmental cues (e.g., the shifting light and temperature conditions on land) and physical stimuli (e.g., polar growth under a lack of buoyancy).

More than one-third of the total detected TAP family gains across the species tree were identified within the streptophyte algae. These gains were probably significant for enabling the water-to-land transition of plant life. In the embryophytes several more TAP families were gained, but dominantly expansions of existing TAP families occurred: More than half (77 out of 135) expansions were detected in embryophytes, confirming a massive expansion of many TAP families concomitant with terrestrialization and subsequent diversification of land plants. The TAP complements of streptophyte algae belonging to the ZCC grade were identified to be most similar to that of embryophytes, consistent with the fact that the ZCC grade and, in particular, the Zygnematophyceae are the closest relatives of land plants (Cheng et al., 2019; Leebens-Mack et al., 2019). In *P. margaritaceum* and *S. muscicola*, secondary expansions of the total TAP complement have occurred, in form of massive small-scale duplications in *P. margaritaceum* and a genome triplication event in *S. muscicola,* resulting in comparable total TAP numbers as in embryophytes. This suggests that a secondary reduction in morphological complexity of an organism does not necessarily lead to a reduction of its TAP family complement (although such a reduction caused by an evolutionary bottleneck has been suggested in other cases, like red algae) (Collén et al., 2013; Petroll et al., 2021). Interestingly, within the ZCC grade, *Z. circumcarinatum,* with the lowest total TAP number in this grade, exhibits the highest concordance of the TAP families presence/absence pattern of embryophytes.

By defining new TAP (sub)families in TAPscan, nine additional (sub)families were detected to be gained in the MRCA of Streptophyta or within streptophyte algae. In previous analyses, TAPs classified as plant specific were dated back to the origin of Streptophyta and hence were considered important for the conquest of land (Wilhelmsson et al., 2017). By defining new subfamilies, we were able to define additional TAP family gains (ALOG, C1HDZ, C2H2-IDD, HD-LD, HD-NDX, HD-SAWADEE, LOB2, MYB-4R and NLP) which might be relevant for enabling terrestrialization (Figure 7).

In conclusion, TAPscan is a powerful tool to comprehensively annotate TAPs with high accuracy. This methodology is now available to the community to foster future analyses. With the ever-increasing number of genomes becoming available, exciting discoveries are expected.

## Supplementary Materials

**Table S1.** Species datasets (DS1-3)

**Table S2.** TAP complements of the investigated species

**Table S3.** Gold standard used for the new TAP (sub)families and information about newly added domains

**Table S4.** Results of statistical tests

**Table S5.** Results of asymmetric Wagner parsimony and overview of observed evolutionary events in previous analyses and literature

**Table S6.** The annotated TAP (sub)families and sequence IDs for *Arabidopsis thaliana* and sensitivity and specificity of the families C1HDZ, C2H2-IDD and ET

**File S1.** TAPscan profile HMMs

**File S2.** Alignments used to generate TAPscan custom profile HMMs

**File S3.** TAPscan classification rules

**File S4.** TAPscan coverage values

**File S5.** Phylogenies of the LBD and HDZ families to support the definition of subfamilies

**File S6.** Pruned phylogeny used as species tree for asymmetric Wagner parsimony analysis

## Data statement

All data is available within the supplementary material (Table S1-S6 and File S1-S6). Additionally, detailed information and access to the resources can be found in the following GitHub repositories:

TAPscan-website:

– URL: https://github.com/Rensing-Lab/TAPscan-v4-website

– This repository contains all the code used to run the TAPscan v4 website, as well as all of the data in its database (TAPs, rules, species list, TAP information, domain list).

Detailed installation instructions are included.

– GPLv3 license

TAPscan-classify standalone tool:

– URL: https://github.com/Rensing-Lab/TAPscan-classify

– This contains the TAPscan classification tool. This consists of a perl script that runs on HMMER output, and assigns TAP families to sequences. The output of this tool is used in the TAPscan v4 website.

– GPLv3 license

TAPscan-classify Galaxy tool:

– GalaxyEU URL: https://usegalaxy.eu/tool_runner?tool_id=toolshed.g2.bx.psu.edu%2Frepos%2Fbgruening%2Ftapscan%2Ftapscan_classify%2F4.74%2Bgalaxy0

– Source code URL: https://github.com/bgruening/galaxytools/tree/master/tools/tapscan

– This is tool is the same as the standalone tool, but provides a wrapper for integration into the Galaxy platform.

– MIT license for the Galaxy wrapper (GPLv3 license for the underlying TAPscan tool) MAdLandDB/Genome Zoo:

– URL: https://github.com/Rensing-Lab/Genome-Zoo

– This repository contains all the sequence fasta files for all species in the Genome Zoo, and therefore all species shown in the TAPscan v4 website. A table with detailed information about each species and the origin of the data is also included

– CC-BY 4.0 license

The TAPscan v4 web presence is available at http://tapscan.plantcode.cup.uni-freiburg.de/ and TAPscan and Genome Zoo can be utilized via https://usegalaxy.eu/.

## Acknowledgments

We thank Stephane Rombauts, Bert De Rybel and Jim Renema from Ghent University, Belgium to provide the unpublished sequence data for *Atrichum angustatum.* The full statement regarding the release of the sequence data for *A. angustatum* can be found in the Genome Zoo Github repository (https://github.com/Rensing-Lab/Genome-Zoo) and the Table S1 (DS3).

