## Supplementary material for "Enhanced sensitivity of TAPscan v4 enables comprehensive analysis of streptophyte transcription factor evolution": File S5

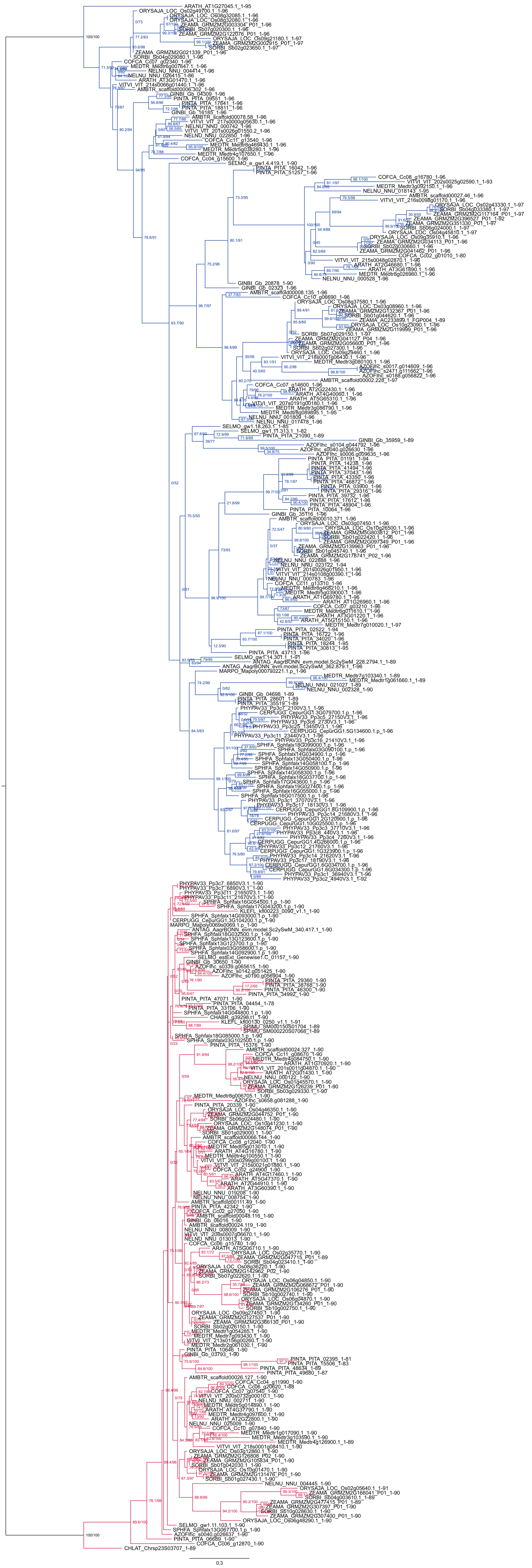

**File S5A. Maximum lieklihood tree of the TAP family HDZ.** This phylogeny was calculated to ensure accurate separation between the subfamilies C1HDZ and C2HDZ. The rooted tree was calculated using the model JTT+I+G4. Included are 335 sequences from species of DS1 belonging to the HDZ subfamilies class I and class II. The clusters of C1HDZ and C2HDZ sequences are colored in blue and red, respectively. The species names are abbreviated in a five-letter code. The numbers at the nodes are SH-aLRT and ultrafast bootstrap support values.

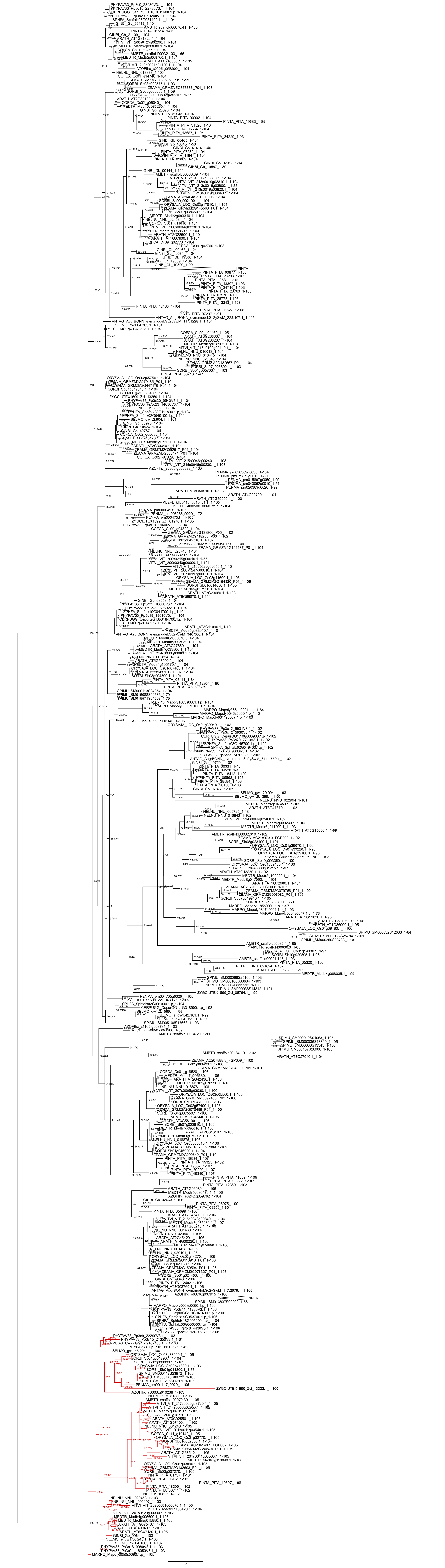

**File S5B. Maximum lieklihood tree of the TAP subfamilies LOB1 and LOB2.** This phylogeny was calculated to ensure accurate separation between the subfamilies LOB1 and LOB2. The rooted tree was calculated using the model JTT+R5. Included are 465 sequences from species of DS1 belonging to the LBD subfamilies LOB1 and LOB2. The cluster of LOB2 is colored in red. The species names are abbreviated in a five-letter code. Clusters in which only sequences belonging to *Pinus taeda* are present have been grouped for clarity. The numbers at the nodes are SH-aLRT and ultrafast bootstrap support values.
